# DNA-utilization loci enable exogenous DNA metabolism in gut Bacteroidales

**DOI:** 10.1101/2025.03.18.642099

**Authors:** Deepti Sharan, Agnieszka Nurek, Joshua Stemczynski, Kristof Turan, Michael J. Coyne, Mary McMillin, Ashley M. Sidebottom, Laurie E. Comstock, Samuel H. Light

## Abstract

The human gut microbiome plays a central role in nutrient metabolism, yet the fate of exogenous nucleic acids within this ecosystem remains poorly understood. Here, we show that multiple Bacteroidales species efficiently metabolize exogenous DNA, with *Bacteroides thetaiotaomicron* converting it into the deaminated nucleobases uracil and xanthine. Using genetic and biochemical approaches, we identify *ddbABCDEF*, a six-gene locus encoding secreted nucleases and an outer membrane transporter, essential for exogenous DNA metabolism in *B. thetaiotaomicron*. Colonization of gnotobiotic mice with *ddbABCDEF* mutants reveals that this pathway significantly alters nucleobase pools in the gut. Comparative genomics demonstrate that *ddbABCDEF* is evolutionarily related to a natural transformation system present in Bacteroidota and has diversified into four distinct subtypes, each linked to unique DNA-processing activities in closely related gut Bacteroidales strains. These findings thus establish DNA as a metabolic substrate in the gut microbiome and reveal a distinctive pathway for nucleobase production with implications for host-microbe interactions.

**SIGNIFICANCE STATEMENT:** The gut microbiome plays a crucial role in nutrient metabolism, yet the fate of extracellular DNA within this ecosystem remains poorly understood. This study identifies *Bacteroidales* species that actively metabolize extracellular DNA, revealing a conserved pathway that converts DNA-derived nucleotides into deaminated nucleobases. We show that *Bacteroides thetaiotaomicron* utilizes a specialized genetic locus, *ddbABCDEF*, to facilitate this process, influencing nucleobase availability in the gut. Comparative genomic analyses suggest that *ddbABCDEF* is evolutionarily linked to bacterial natural transformation systems but has diverged into distinct metabolic subtypes. These findings establish DNA as a metabolic substrate in the gut microbiome, with potential implications for microbial ecology, host-microbe interactions, and gut health.

## INTRODUCTION

The human gut microbiome comprises a complex community of microorganisms that play pivotal roles in host physiology. A key mechanism by which the gut microbiome impacts host health is through the metabolism of dietary- and host-derived compounds, products of which include bioactive metabolites that influence systemic functions (1). While microbial metabolism of carbohydrates and proteins has been extensively studied, the processing of dietary nucleic acids by the gut microbiome remains underexplored.

Nucleic acids are abundant in plant- and animal-derived foods and undergo partial enzymatic digestion in the host gastrointestinal tract, yielding nucleotides, nucleosides, and nucleobases (2). While these molecules are partially absorbed in the small intestine, some fraction reaches the lower gastrointestinal tract, where the microbial density is highest (3, 4). Additionally, host-derived nucleic acids, including DNA from sloughed epithelial cells, infiltrating immune cells, and lysed microbial cells, further contribute to the pool of exogenous nucleic acids available for microbial metabolism (5, 6). The presence of dietary and host-derived DNA in fecal matter confirms that nucleic acids remain a persistent resource available for microbial breakdown in the gut (3, 4, 7, 8).

Recent studies have implicated nucleic acids as an important resource for the gut microbiome. For instance, ribose – the sugar backbone of RNA – has been shown to provide a growth substrate for *Bacteroides thetaiotaomicron*, a prominent gut symbiont, within the gut (9). Additionally, microbial metabolism of nucleic acids may contribute bioactive metabolites with systemic effects. For example, the microbiome-derived nucleosides inosine and guanosine enhance immune responses, improving outcomes in cancer immunotherapy and tolerance against food allergens, respectively (10, 11). Likewise, nucleobases such as xanthine and hypoxanthine have been shown to promote intestinal wound healing and reinforce mucosal barrier function (12). Multiple lines of evidence thus suggest that microbial nucleic acid metabolism may have significant implications for microbial ecology and host biology within the gut.

In this study, we investigate mechanisms of exogenous DNA metabolism by gut bacteria. Through genetic, biochemical, and metabolomic approaches, we demonstrate that *B. thetaiotaomicron* utilizes a locus that we name *ddbABCDEF* to mediate exogenous DNA degradation and the conversion of DNA nucleotides into deaminated nucleobases, such as xanthine and uracil. Comparative genomic analyses reveal that the *ddbABCDEF* locus is part of a larger family of functionally diverse nucleic acid processing systems widespread among gut Bacteroidales, with distinct subtypes linked to unique DNA-depleting activities. These findings define a novel class of DNA metabolism systems in the gut microbiome and offer new insights into the ecological and metabolic roles of microbial DNA processing in the mammalian gut.

## RESULTS

### Human fecal microbial communities and gut *Bacteroides* deplete exogenous DNA

To assess whether human gut microbial communities metabolize exogenous DNA, we inoculated anaerobic human fecal cultures collected from five different donors with herring sperm DNA or synthetic oligonucleotides and monitored DNA depletion over time. DNA levels declined progressively in all fecal cultures tested, but the rate of depletion varied considerably between donors (**Figure 1A & 1B, Figure S1**). These results demonstrate that the gut microbiome can effectively degrade exogenous nucleic acids, with potential donor-dependent distinctions in the nature of this activity.

**Figure 1.**
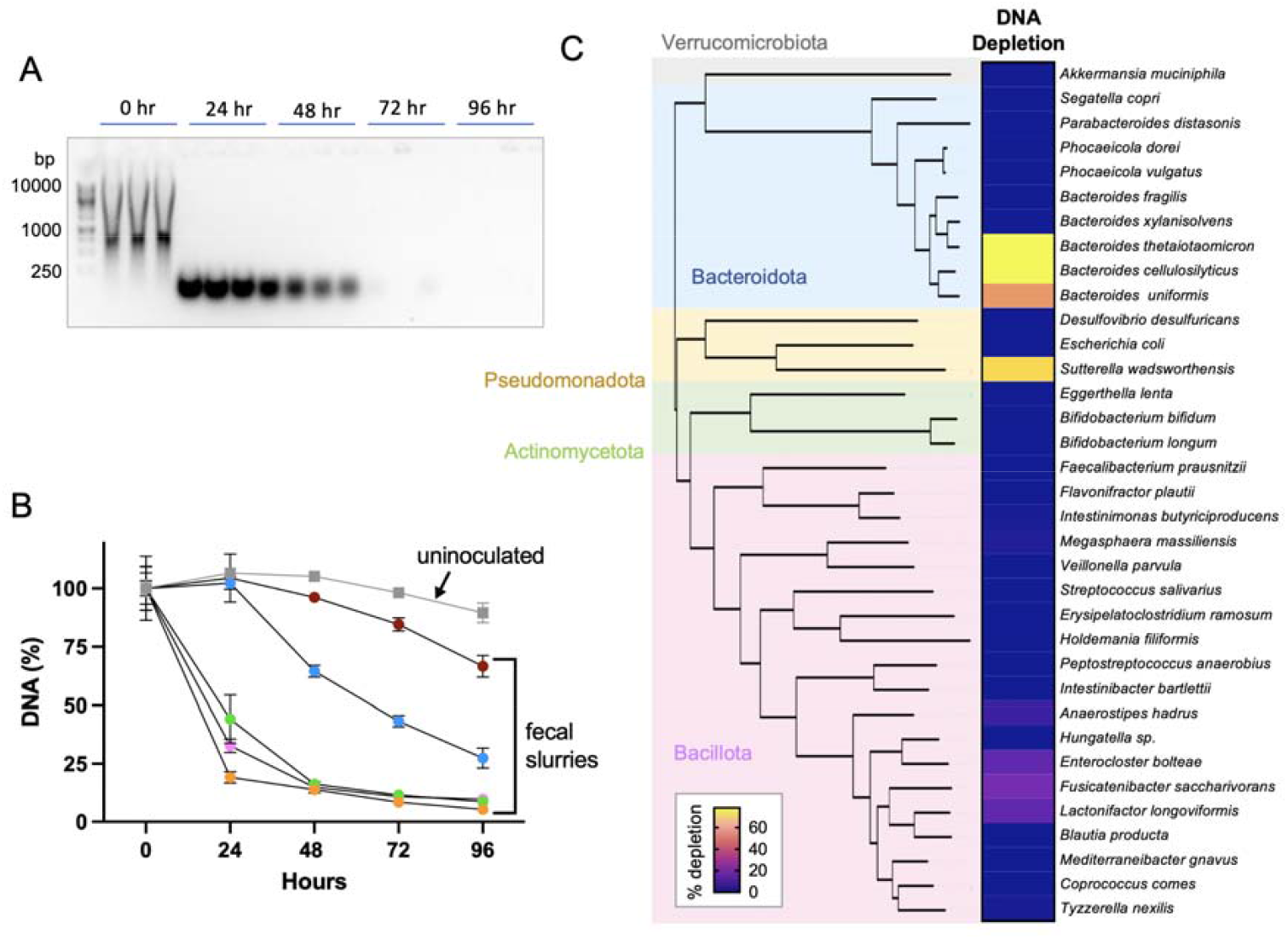
Human fecal microbial communities and gut Bacteroidales deplete exogenous DNA. (A) Representative agarose gel showing progressive degradation of herring sperm DNA over time in a human fecal slurry (donor 1), demonstrating microbial DNA depletion activity. (B) Quantification of DNA depletion in fecal slurries from five different donors over time. (C) DNA depletion activity of monocultures from taxonomically diverse gut bacterial strains.

To identify specific microbial taxa responsible for the observed phenotype, we next screened a panel of 35 taxonomically diverse gut bacterial strains for DNA-depleting activity. Among tested strains, those belonging to four species—*Bacteroides cellulosilyticus, Bacteroides thetaiotaomicron, Bacteroides uniformis*, and *Sutterella wadsworthensis*— exhibited robust DNA-depleting activity (**Figure 1C, Figure S2**). These results thus show that exogenous DNA is depleted by the gut microbiome and implicate the gut Bacteroidales as playing a significant role in this activity.

### *B. thetaiotaomicron ddbABCDEF* is essential for exogenous DNA depletion

Bacteroidales cultures incubated with herring sperm DNA revealed a gradual decrease in fragment size over time, consistent with extracellular nuclease activity (**Figure S1**). Given that secreted nucleases are commonly involved in DNA metabolism, we tested whether *B. thetaiotaomicron* produced such enzymes. Consistent with secreted factors contributing to DNA degradation, we found that culture supernatants from *B. thetaiotaomicron* strain DFI.1.47 exhibited robust nuclease activity (**Figure 2A**).

**Figure 2.**
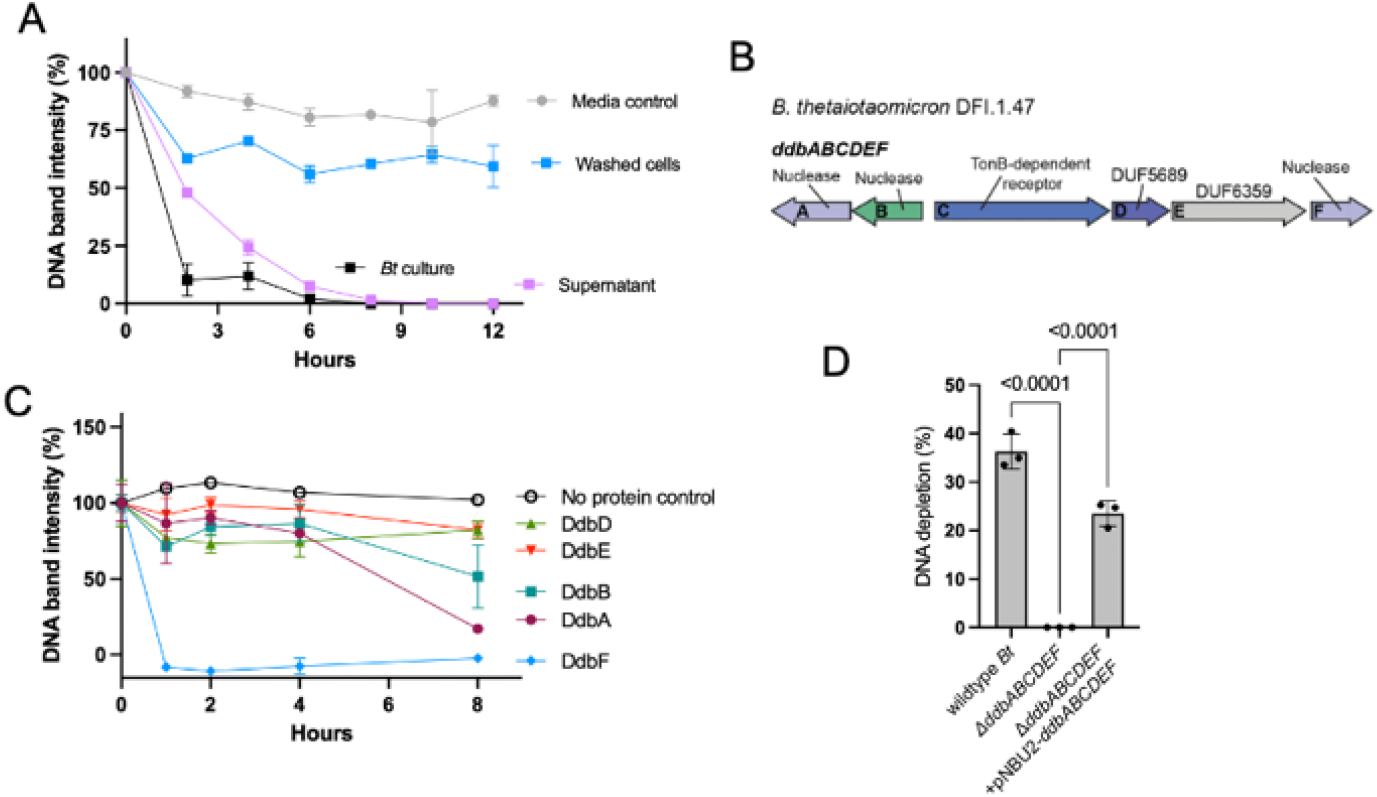
The *ddbABCDEF* locus encodes secreted nucleases and is essential for DNA depletion in *B. thetaiotaomicron*. (A) DNA depletion activity of *B. thetaiotaomicron* cultures, comparing whole cells and secreted factors. (B) The *ddbABCDEF* locus. DUF stands for domain of unknown function. (C) DNA degradation activity of purified recombinant proteins encoded by the *ddbABCDEF* locus. (D) DNA depletion activity of wild-type *B. thetaiotaomicron* compared to uncomplemented and complemented Δ*ddbABCDEF* mutants. Statistical significance was assessed using one-way ANOVA followed by Dunnett’s post hoc test

To identify the genetic basis for this activity, we searched the *B. thetaiotaomicron* DFI.1.47 genome for putative nucleases with predicted extracellular or periplasmic localization. This analysis led to the identification of a six-gene locus, which we designated *ddbABCDEF* (for *<u>D</u>*NA *<u>D</u>*epletion in *<u>B</u>*acteria). The locus encodes three predicted nucleases (DdbA, DdbB, and DdbF) – including a homolog of a previously identified variant (*ddbF*) proposed to encode an extracellular nuclease required for bile-dependent biofilm formation (13) – alongside an outer membrane transporter (DdbC), a hypothetical protein (DdbD), and an additional predicted extracellular protein (DdbE) (**Figure 2B**). Consistent with the annotated functions, heterologous expression and *in vitro* nuclease assays confirmed that DdbA, DdbB, and DdbF exhibit DNA-degrading activity (**Figure 2C**).

In gut Bacteroidales, distinct genetic loci encode proteins with degradative enzymes and outer membrane transporters that facilitate uptake of specific complex exogenous polymers, including polypeptides and polysaccharides (14–16). Considering these parallels, we reasoned the *ddbABCDEF* locus might be relevant for DNA utilization *B. thetaiotaomicron* DFI.1.47. To test this hypothesis, we generated and assayed a scarless deletion mutant (Δ*ddbABCDEF*) and a complemented (Δ*ddbABCDEF* + pNBU2-*ddbABCDEF*) strain. We found the Δ*ddbABCDEF* strain was completely deficient in DNA depletion activity, but that this defect was restored by complementation (**Figure 2D**). These results establish that *ddbABCDEF* is required for exogenous DNA degradation in *B. thetaiotaomicron* DFI.1.47.

### *ddbABCDEF* is required for deaminated nucleobase production from DNA

We next sought to investigate the physiological significance of *B. thetaiotaomicron* DNA metabolism. As preliminary experiments revealed that exogenous DNA provided a poor sole carbon, nitrogen, or phosphorus source, we asked whether *B. thetaiotaomicron* could metabolize DNA into biologically relevant secondary products. We performed targeted metabolomics analyses of *B. thetaiotaomicron* culture supernatants grown in the presence or absence of exogenous DNA and observed significantly elevated levels of the deaminated nucleobases xanthine and uracil in cultures supplemented with DNA (**Figure 3A**).

**Figure 3.**
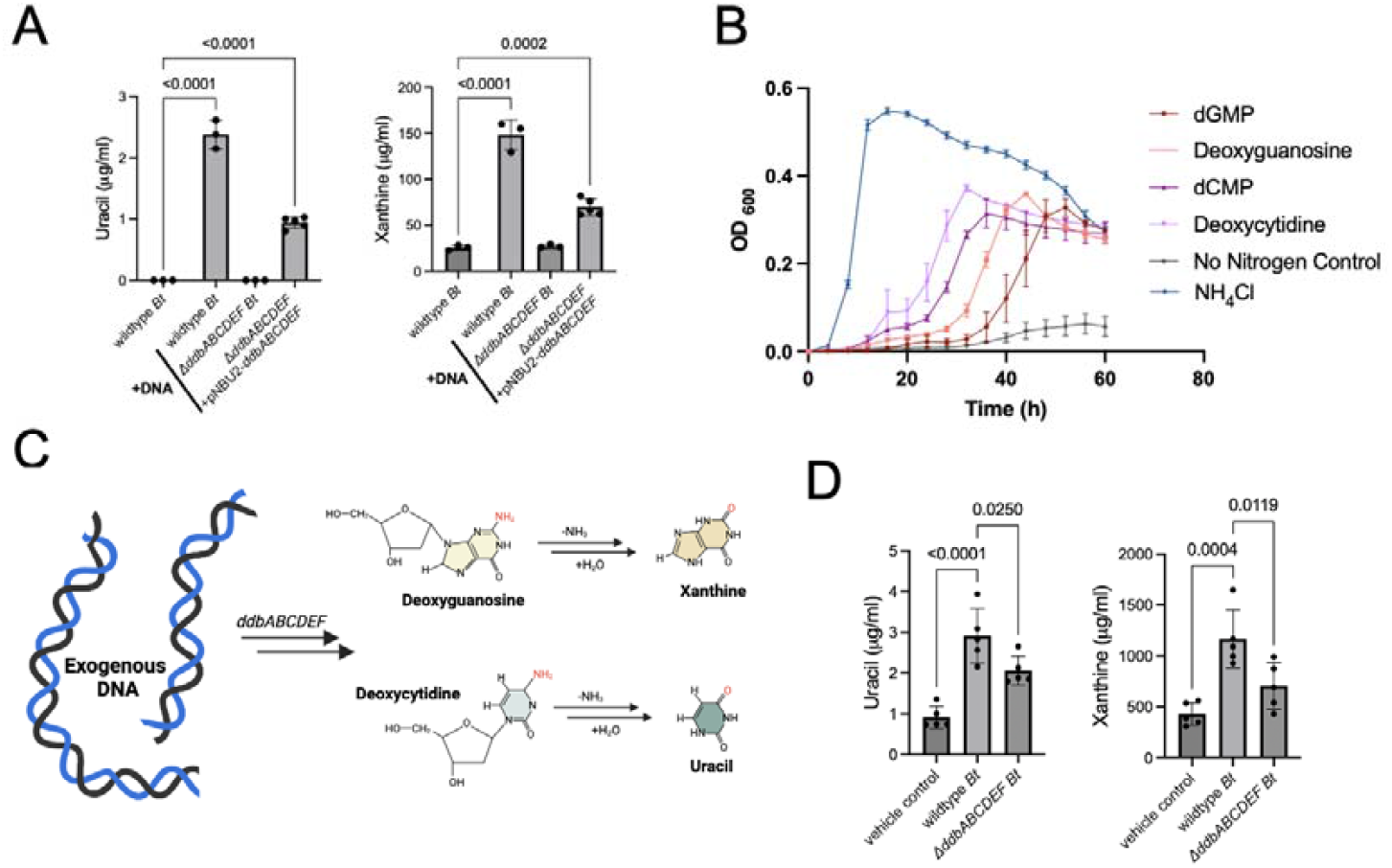
Exogenous DNA fuels production of deaminated nucleobases by *B. thetaiotaomicron*. (A) Quantification of uracil and xanthine levels in culture supernatants of *B. thetaiotaomicron* grown with or without exogenous herring sperm DNA. (B) Growth of *B. thetaiotaomicron* in chemically defined media supplemented with different nitrogen sources. (C) Proposed pathway linking DNA degradation to nucleobase production. (D) Uracil and xanthine levels in the cecal contents of germfree mice gavaged with indicated *B. thetaiotaomicron* strains or a vehicle control. Statistical significance was assessed using one-way ANOVA followed by Dunnett’s post hoc test

To clarify the pathway responsible for this conversion, we systematically tested the ability of *B. thetaiotaomicron* to metabolize DNA-derived nucleotides, nucleosides, and nucleobases. We found that cytosine and guanine nucleotides could serve as sole nitrogen sources, albeit suboptimally, and were readily converted into uracil and xanthine, respectively (**Figure 3A & 3B, Figure S3**). Notably, deletion of *ddbABCDEF* abolished uracil and xanthine production from exogenous DNA but had no effect on their generation from deoxycytidine and deoxyguanosine nucleosides (**Figure 3A, Figure S4**). This finding indicates that *ddbABCDEF* is specifically required for exogenous DNA uptake and that other presently unidentified downstream enzymes are required for subsequent reactions to generated deaminated nucleobases (**Figure 3C)**.

### *ddbABCDEF* promotes deaminated nucleobase production in the mouse gut

To test whether *ddbABCDEF*-mediated DNA metabolism contributes to nucleobase production *in vivo*, we colonized gnotobiotic mice with wildtype or Δ*ddbABCDEF* mutant *B. thetaiotaomicron* strains and quantified uracil and xanthine levels in the cecum. Monocolonization with wildtype *B. thetaiotaomicron* resulted in a significant increase in cecal uracil and xanthine concentrations relative to germ-free controls (**Figure 3D**). Compared to wildtype *B. thetaiotaomicron*, Δ*ddbABCDEF* monocolonization resulted in a significant reduction in the cecal concentration of these metabolites (**Figure 3D**). These results provide evidence that *B. thetaiotaomicron* generates deaminated nucleobases in the gut and that *ddbABCDEF* is essential for maximal flux through this pathway.

### *ddbABCDEF* is related to the *Riemerella anatipestifer* natural transformation system

Having identified a role for the *ddbABCDEF* locus in exogenous DNA metabolism, we next sought to investigate the functions of non-nuclease proteins encoded within the locus. Natural transformation systems enable bacteria to import and integrate foreign genetic material, thereby serving as a mechanism for horizontal gene transfer. We observed that DdbC and DdbD share homology with proteins RA0C_RS04915 and RA0C_RS04920, which were recently identified as essential components of the natural transformation machinery in the bird pathogen *Riemerella anatipestifer* and were proposed to facilitate DNA transport across the outer membrane in this Bacteroidota species (17). Further examination revealed that these *R. anatipestifer* genes neighbored a gene with sequence homology to the *ddbB* nuclease, thus presenting a locus strikingly similar to *ddbBCD* in *B. thetaiotaomicron* (**Figure 4A**).

**Fig 4.**
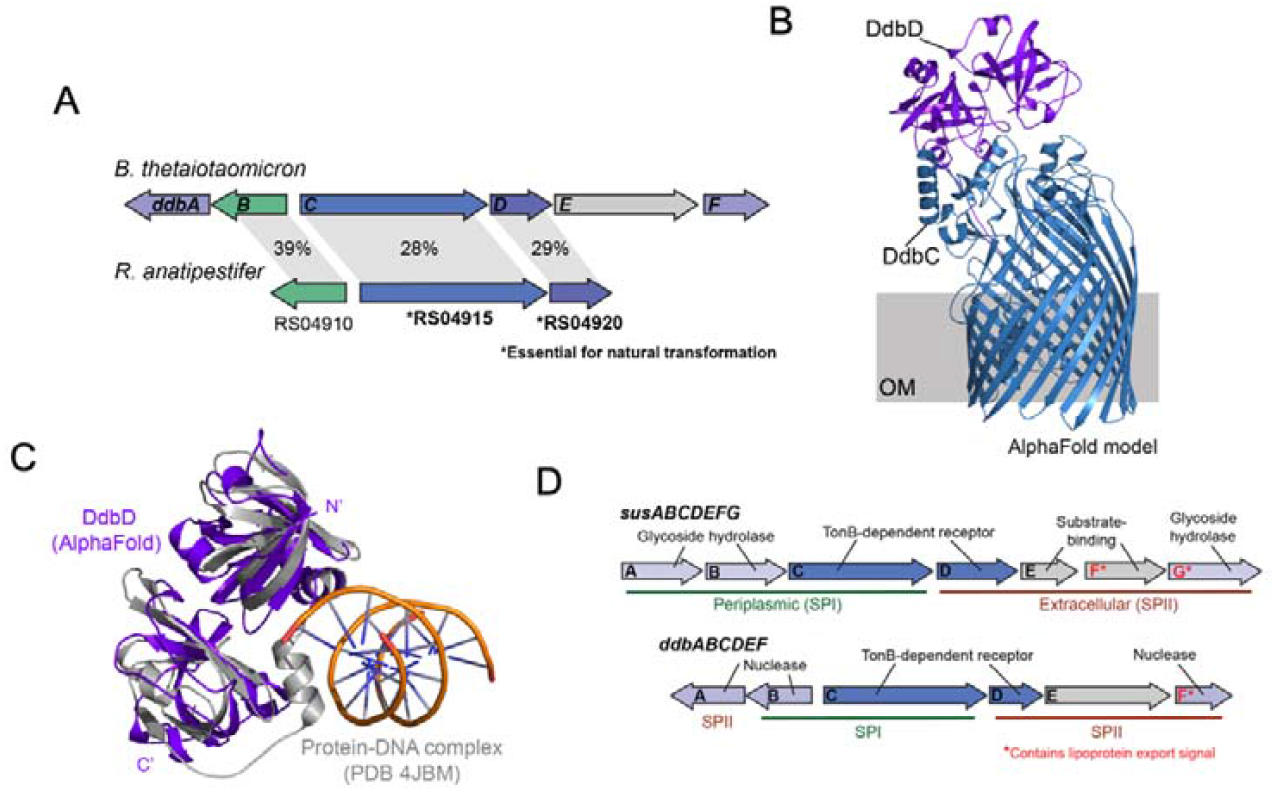
The *ddbABCDEF* locus resembles both natural transformation and polysaccharide-utilization loci. (A) Schematic comparison of the *ddbABCDEF* locus with a homologous gene cluster in *Riemerella anatipestifer* involved in natural transformation. Percentages indicate amino acid sequence identity between homologous proteins. Names refer to locus tags RA0C_RS049*XX*. (B) AlphaFold-predicted structural model of the DdbC-DdbD complex. The gray box indicates the approximate position of the outer membrane (OM). (C) Structural alignment of DdbD with the DNA-binding domain of murine p202 complex with double-stranded DNA (PDB: 4JBM). (D) Comparison of the *ddbABCDEF* locus with the *B. thetaiotaomicron* starch-utilization system (*susABCDEFG*), the archetypal polysaccharide-utilization loci. Signal peptide classifications (SPI vs. SPII) indicate the predicted localization of encoded proteins.

The conserved synteny of *R. anatipestifer* natural transformation genes with DdbC and DdbD led us to interrogate the relationship between these proteins. DdbC is annotated as a transmembrane subunit of a TonB-dependent outer membrane transporter, while DdbD is annotated as a domain of unknown function (DUF5689). TonB-dependent transporters typically include an extracellular substrate-binding subunit encoded by a gene directly adjacent to the transmembrane subunit (16, 18). Based on this organization, we hypothesized that DdbD may act as the substrate-binding partner for DdbC.

AlphaFold protein structural modeling was recently shown to enable detection of interactions between *Bacteroides* TonB-dependent transporter transmembrane and substrate-binding subunits (19). Consistent with DdbC and DdbD forming a heterodimeric complex, AlphaFold3 predicted a high confidence DdbC/DdbD interface that reasonably positioned DdbD for a potential substrate-binding role (**Figure 4B**). Despite possessing low levels of sequence identity, we further observed that DdbD AlphaFold structure exhibited high structural similarity to previously characterized DNA-binding proteins, including the DNA-binding domain of p202, a murine immune protein that non-specifically binds double-stranded DNA (20) (**Figure 4C**). Collectively, these observations are consistent with DdbCD being part of a structurally distinctive family of outer membrane transporters associated with natural transformation and DNA metabolism in bacteria.

### *ddbABCDEF* resembles gut Bacteroidales polysaccharide-utilization loci

Numerous polysaccharide-utilization loci (PULs) that encode proteins responsible for metabolism of a specific polysaccharides have been identified in gut Bacteroidales (reviewed in (14)). PULs typically encode (a) extracellular polysaccharide-binding proteins that position the polysaccharide for hydrolysis, (b) extracellular glycoside hydrolases that hydrolyze polysaccharides to oligosaccharides, (c) a TonB-dependent outer membrane transporter that transports oligosaccharides into the periplasm, and (d) periplasmic glycoside hydrolases that hydrolyze oligosaccharides into mono- and disaccharides.

Within PULs, the distinct cellular localization of proteins is often reflected in the signal peptides that govern their secretion. Periplasmic proteins generally contain an SPI signal peptide that results in retention in the periplasm, whereas extracellular proteins typically contain an SPII signal peptide that results in lipid-mediated anchoring to the outer membrane (14). Lipoprotein export signals further mark a subset of extracellular proteins for packaging into secreted outer membrane vesicles (21–23).

As illustrated by a comparison to the emblematic starch PUL *susABCDEFG*, the *ddbABCDEF* loci is notable for its PUL-like organization, just differing in the nature of the polymer-hydrolyzing enzymes (**Figure 4D**). DdbA and DdbF have SPII signal peptides and hydrolyze nucleic acids, analogous to the extracellular polysaccharide hydrolase SusG. DdbCD is a TonB-dependent transporter that likely transports products generated from extracellular enzymes into the periplasm, analogous to SusCD. DdbB has a SPI signal peptide and hydrolyzes nucleic acids, analogous to periplasmic oligosaccharide hydrolases SusA and SusB. Finally, DdbE has a SPII and encodes a domain of unknown function that could engage in substrate-binding properties, analogous to SusEF. Collectively, these analyses thus demonstrate parallels between *ddbABCDE*F and both natural transformation and polymer-degrading Bacteroidales loci.

### Gut Bacteroidales encode multiple *ddb* subtypes

To assess the distribution of *ddbABCDEF*-like loci, we performed a comparative genomic analysis across diverse members of the phylum Bacteroidota. Consistent with the DNA-depleting activity observed for multiple Bacteroidales strains in our initial screen (**Figure 1C**), bioinformatics analyses identified numerous loci across multiple species anchored by the presence of a DdbC transporter homolog (**Figure 5A**). These loci consistently encoded annotated secreted nucleases, suggesting a conserved relevance for exogenous nucleic acids (**Figure 5A**).

**Figure 5.**
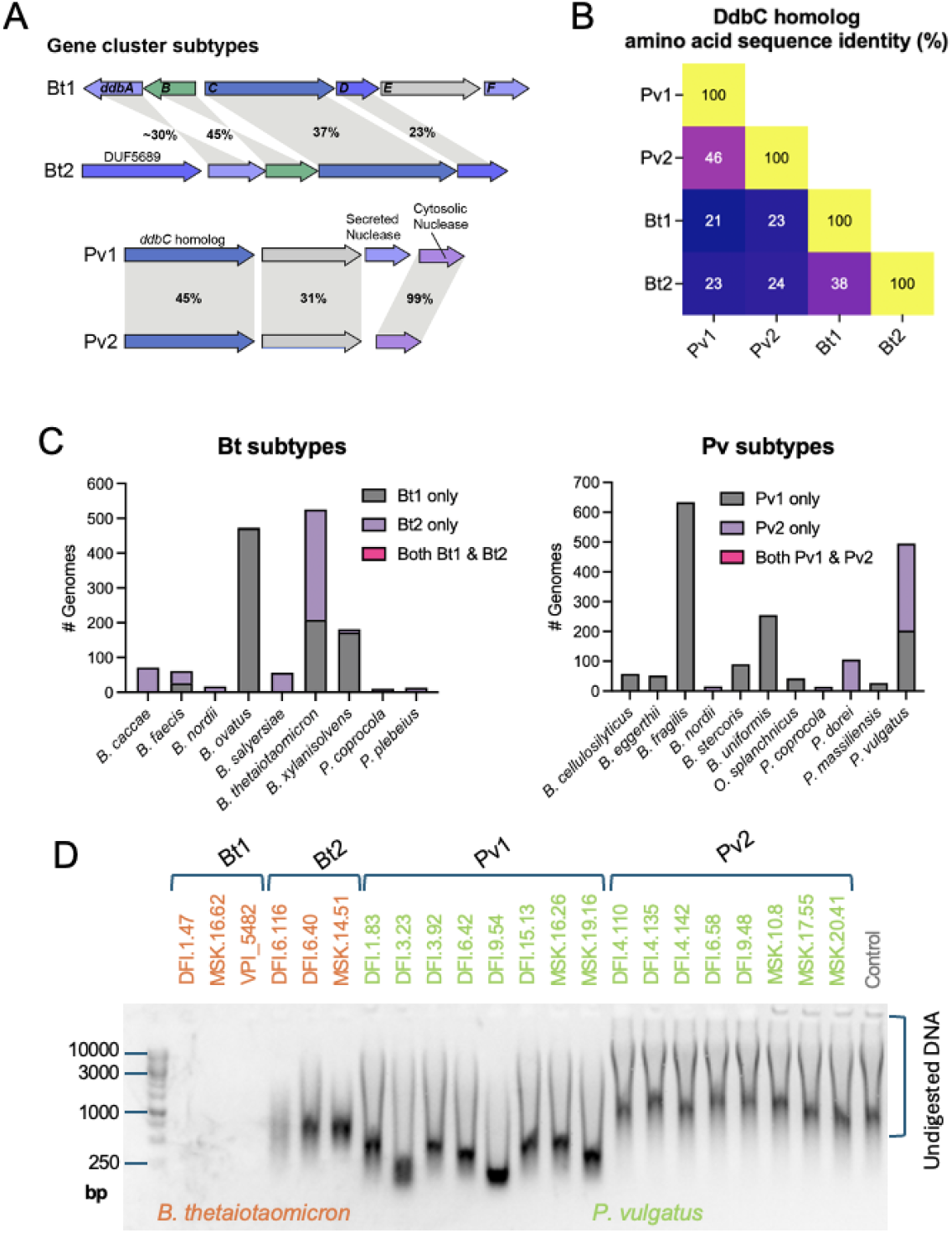
Distinct *ddbABCDEF* -like subtypes are present in gut Bacteroidales and confer variable DNA-depleting activities. (A) Representative *ddbABCDEF* -like subtype gene clusters from *B. thetaiotaomicron* DFI.1.47 (Bt1) *B. thetaiotaomicron* MSK.14.51 (Bt2) and *P. vulgatus* DFI.1.83 (Pv1) and *P. vulgatus* DFI.4.110 (Pv2). Percentages indicate amino acid sequence identity between homologous proteins. (B) Heatmap illustrating amino acid sequence identity among DdbC homologs from Bt1, Bt2, Pv1, and Pv2 *ddbABCDEF*-like gene clusters. (C) Distribution of *ddbABCDEF*-like subtypes across gut Bacteroidales genomes. (D) Activity of Bacteroides and Phocaeicola strains encoding different *ddbABCDEF* subtypes incubated with herring sperm DNA.

Further analyses revealed that *ddbABCDEF*-like loci could be subdivided into two types based on DdbC protein sequence homology. DdbC homologs from *B. thetaiotaomicron* and *Phocaeicola vulgatus* exemplify these two types and share approximately 25% sequence identity. *B. thetaiotaomicron* and *P. vulgatus* types could be further parsed into subtypes that share less than 50% sequence identity, which we designate as Bt1 and Bt2 (for *B. thetaiotaomicron* 1 and 2) and Pv1 and Pv2 (for *P. vulgatus* 1 and 2) subtypes (**Figure 5B**).

The distribution of *ddb* subtypes did not generally track with taxonomy and exhibited a striking pattern of mutual exclusivity across genomes. While some genomes encoded both Bt- and Pv-type loci, different subtypes (such as Bt1 and Bt2 or Pv1 and Pv2) were rarely found in the same genome (**Table S1**). This exclusivity was especially apparent when comparing strains of a single species. Among 525 *B. thetaiotaomicron* genomes analyzed, 208 encoded Bt1 and 317 encoded Bt2, but none encoded both (**Figure 5C, Table S1**). Similarly, among 495 *P. vulgatus* genomes analyzed, 203 encoded Pv1 and 292 encoded Pv2, but none encoded both (**Figure 5C, Table S1**). We further identified five additional species with comparable patterns of variable strain-level mutual exclusivity across multiple subtypes (**Figure 5C, Table S1**).

### *ddb* subtypes are associated with distinct DNA-depleting activities in closely related gut Bacteroidales

The presence of multiple *ddb* subtypes across *gut Bacteroidales raises* the possibility that these variants encode functionally distinct DNA-processing activities. To assess this, we evaluated the DNA-depleting capacity of *B. thetaiotaomicron* and *P. vulgatus* strains carrying different *ddb* subtypes. Across tested strains, those encoding the same subtype exhibited consistent DNA degradation phenotypes, but striking differences emerged between subtypes. Strains harboring the Bt1 subtype fully degraded exogenous DNA, while Bt2 strains exhibited only partial degradation (**Figure 5D**). Similarly, Pv1 strains partially depleted DNA, whereas Pv2 strains exhibited minimal DNA-depleting activity (**Figure 5D**). These results suggest that *ddb* subtypes have diverged functionally, with some variants more effective at degrading extracellular DNA than others.

## DISCUSSION

This study highlights the role of *B. thetaiotaomicron* and related gut Bacteroidales in the metabolism of exogenous DNA. Our findings demonstrate that the *ddbABCDEF* locus encodes multiple nucleases and a predicted TonB-dependent transporter and is essential for DNA degradation and deaminated nucleobase production in the mouse gut. These components presumably operate in concert to mediate exogenous DNA cleavage and transport, enabling *B. thetaiotaomicron* to metabolize DNA to deaminated nucleobases.

A particularly intriguing feature of the *ddbABCDEF* locus is its evolutionary and functional resemblance to the natural transformation machinery of *R. anatipestifer*, a member of the Flavobacteriales within the Bacteroidota phylum. Unlike Flavobacteriales, gut Bacteroidales are not known to be naturally competent. Consistent with this, our attempts to detect DNA uptake in a diverse set of gut *Bacteroidales* yielded inconsistent results. This suggests that while *ddbABCDEF* may share a common ancestral origin with natural transformation systems, it has since diverged functionally.

Notably, a comparable functional versatility exists in *Pseudomonadota*, where a pilus-based apparatus facilitates DNA transport across the outer membrane (24). In *Pseudomonadota*, this machinery is used for either natural transformation or DNA-based nutrient acquisition, depending on species-specific adaptations and environmental context (24–26). The characterization of *ddbABCDEF* thus contributes to a growing body of evidence indicating that phylogenetically diverse bacteria have evolved distinct yet functionally analogous mechanisms to acquire and metabolize extracellular DNA.

Discovery of multiple *ddbABCDEF* subtypes further underscores the functional diversity of DNA processing systems in gut Bacteriodota. The observed mutual exclusivity of subtypes across strains and the association of this with distinct DNA-depleting activities strongly suggests that unknown factors functionally differentiate *ddbABCDEF* subtypes. These findings thus provide evidence that significant gaps remain in our understanding of the full functional potential of exogenous DNA uptake and processing by gut Bacteriodota, which could be an interesting topic for future study.

Finally, our findings also raise questions about the ecological and host-relevant impacts of microbial DNA metabolism. The production of deaminated nucleobases, such as xanthine and uracil, within the mouse gastrointestinal tract suggests potential downstream effects on host physiology. These metabolites may act as signaling molecules, modulate immune responses, or alter gut barrier function. Prior studies have shown that nucleobase-derived metabolites influence systemic processes, including immune regulation and gut barrier regeneration. While these molecules could be generated via multiple routes, our findings establish the microbial metabolism of exogenous DNA as a plausible source of bioactive nucleobase-derived metabolites worthy of future interrogation.

## MATERIALS AND METHODS

### Bacterial strains and growth conditions

Bacterial strains used in this study were acquired from either the American Type Culture Collection (ATCC, Manassas, Virginia) or the Duchossois Family Institute (DFI) Biobank strain collection (University of Chicago, Illinois) and are listed in **Table S2** (27, 28). Strains were routinely cultured under anaerobic conditions in a Coy vinyl anaerobic chamber (2-5% H_2_, 2-5% CO_2_, N_2_ balance) at 37°C. Growth media included Brain Heart Infusion Medium with Hemin (BHIM), basal, and chemically defined media. The chemically defined medium contained 1 g/L NH_4_Cl, 6 g/L Na_2_HPO_4_, 3 g/L KH_2_PO_4_, 0.5 g/L NaCl, 14.7 mg/L CaCl_2_, 246 mg/L MgSO_4_·7H_2_O, 0.5% glucose, 0.05% L-cysteine, 2.5 mg/L vitamin K3, 2 mg/L FeSO_4_, 5 mg/L hemin, 5 ng/mL vitamin B12. Basal medium was formulated as previously described and contained 19.2 g/L Na_2_HPO_4_·7H_2_O, 4.5 g/L KH_2_PO_4_, 0.75 g/L NaCl, 1.5 g/L NH_4_Cl, 2 g/L sodium acetate, 1.5 g/L sodium formate, 0.1 g/L tryptone, 0.1 g/L Bacto yeast extract, 1 g/L MgSO_4_, and an 1x stock of minerals and vitamins and was adjusted to pH 6.5. The 1x minerals and vitamins stock solution contained 0.1 mg/L FeCl_2_·4H_2_O, 0.846 mg/L MnSO_4_·H_2_O, 0.028 mg/L ZnSO_4_·7H_2_O, 0.148 mg/L CaCl_2_·2H_2_O, 0.002 mg/L CuSO_4_·5H_2_O, 0.002 mg/L CoCl_2_·7H_2_O, 0.002 mg/L H_3_BO_3_, 0.002 mg/L Na_2_MoO_4_·2H_2_O, 50 mg/L NaCl, 1.2 mg/L tri-sodium citrate, 0.05 mg/L biotin, 0.1 mg/L D-pantothenic acid, 0.05 mg/L lipoic acid, 0.1 mg/L niacinamide, 0.1 mg/L para-aminobenzoic acid, 0.1 mg/L pyridoxal HCl, 0.05 mg/L riboflavin, 0.1 mg/L thiamine HCl, and 0.01 mg/L vitamin B12 (29). Anaerobic growth curves were obtained by shaking cells at 200 rpm and measuring the optical density at 600 nm (OD_600_) every 20 minutes for 72 hours, using a LogPhase 600 (BioTek) plate reader.

### Human fecal microbial community DNA depletion assays

Fecal samples from five healthy, antibiotic-free individuals were collected under an IRB-approved protocol (University of Chicago Protocol IRB20-1384). Samples were frozen in anaerobic 20% glycerol and stored at -80°C until use. Thawed samples were washed in phosphate-buffered saline (PBS) and resuspended in basal medium supplemented with 0.5 mg/mL herring sperm DNA (Sigma D3159) or random sequence 60-bp synthetic oligonucleotides (Integrated DNA Technologies). Cultures were incubated anaerobically at 37°C for 96 hours. Cultures were centrifuged at 5000 x *g* for 10 minutes at 4°C, and 10 μL of supernatant was mixed with 6x TriTrack DNA loading dye (Sigma R1161) before loading onto a 1% agarose gel. Electrophoresis was conducted at 120V for 45 minutes using a Bio-Rad Sub-Cell GT system. DNA bands were visualized using an iBright 1500 imaging system, and band intensities were analyzed with ImageJ software (30).

### Bacterial strain DNA depletion assays

Strains grown overnight anaerobically in BHIM were inoculated into basal media supplemented with 0.5 mg/mL DNA and incubated anaerobically at 37°C for 96 hours. DNA depletion was visualized via agarose gel electrophoresis, with band intensities measured in ImageJ software.

### Nitrogen utilization assays

For experiments testing sole nitrogen sources, *B. thetaiotaomicron* DFI.1.47 overnight anaerobic cultures were washed three times in PBS and before inoculation into chemically defined medium in which NH_4_Cl was replaced with 2 mM nucleic acids, nucleotides, nucleosides or nucleobases (from Sigma or Fisher Scientific).

### *B. thetaiotaomicron* genetic manipulation and mutant generation

A scarless deletion mutant (Δ*ddbABCDEF)* was generated in *B. thetaiotaomicron* DFI.1.47 using counter-selection with the *Bacteroides* suicide vector pLGB13 (31). Flanking 1000-bp regions of *ddbABCDEF* were PCR amplified and cloned into pLGB13. The construct was introduced into *B. thetaiotaomicron via* conjugation with *E. coli* S17-λ*pir*. Cointegrants were selected on BHIM plates containing erythromycin (50 μg/mL) and gentamycin (200 μg/mL). Deletion mutants were isolated by counter-selection on anhydrotetracycline (50 ng/mL) and were confirmed by PCR. The entire *ddbABCDEF* operon from *B. thetaiotaomicron* DFI.1.47 was cloned into the expression vector pNBU2 and introduced into the Δ*ddbABCDEF* mutant strain by conjugal transfer for complementation studies (32).

### Heterologous protein expression and purification

His-tagged versions of DdbA, DdbB, DdbD, DdbE, and DdbF were created by cloning each of these genes into pET28a, and transforming chemically-competent *E. coli* BL21 (DE3) for expression as previously described (33, 34). Briefly, cultures were grown to OD_600_ 0.7-1.0, shaking (RPM 200) at 37° C, before induction with 1 mM isopropyl β-D-1-thiogalactopyranoside and overnight incubation at 20°C. After which, cells were resuspended in lysis buffer containing 300 mM NaCl, 1 mM dithiothreitol, 10 mM imidazole, 50 mM Tris-HCl (pH 7.5), and lysozyme (50 mg/mL). Resuspend cells were placed in an ice water bath, subjected to eight 30 second sonication pulses, and then centrifugated at 40,000 x *g* for 30 minutes at 4°C. His-tagged proteins were purified from the resulting lysate using Profinity IMAC resin, washed, and eluted in 500 mM imidazole buffer. Purity was assessed by SDS-PAGE.

### Nuclease assays

Nuclease activity was assessed in bacterial supernatants, cell lysates, or his-tagged purified proteins. *B. thetaiotaomicron* cultures were grown anaerobically at 37°C for 72 hours, centrifuged at 5000 x *g*, and passed through a 0.22 μm membrane filter. DNA substrates (500 ng/mL) were incubated with 100 μL of supernatant, lysate, or 10 μg of purified protein at 37°C. Reactions were analyzed by agarose gel electrophoresis over time (0-12 hours), and band intensity was quantified using ImageJ software.

### Metabolite analysis

Culture supernatants from *B. thetaiotaomicron* strains incubated in chemical defined medium with or without 0.5 mg/mL herring sperm DNA or 0.15 mg/mL deoxycytidine and deoxyguanosine were collected after 72 hours. Uracil and xanthine concentrations were quantified using gas chromatography-mass spectrometry (GC-MS) and liquid chromatography-mass spectrometry (LC-MS), respectively.

### Gnotobiotic mouse experiments

All mouse experiments were approved by the Institutional Animal Care and Use Committee (IACUC) at the University of Chicago and were conducted in accordance with ethical regulations for animal research. Germ-free C57BL/6J mice (5-10 weeks old) were housed in gnotobiotic isolators at the Gnotobiotic Research Animal Facility (GRAF). Female adult mice were orally gavaged with 200 µL of log-phase cultures of either wild-type *B. thetaiotaomicron* DFI.1.47 or the Δ*ddbABCDEF* mutant strain. Cecal contents were harvested on day 7 and subjected to metabolomic analysis to quantify uracil and xanthine concentrations, as described above.

### Protein Structural Modeling

To predict the structure of the DdbC/DdbD complex, AlphaFold3 was used with default parameters (35). Input sequences for DdbC and DdbD were retrieved from *B. thetaiotaomicron* DFI.1.47 and submitted for co-folding analysis. Model confidence was evaluated using the predicted local distance difference test (pLDDT) and predicted aligned error (PAE) scores. Structural alignments were performed using PyMOL (Schrödinger, LLC) and homology comparisons to known DNA-binding proteins were assessed using the DALI server (36).

### Comparative genomics analyses

NCBI accessions of DdbABCDEF proteins are as follows: WP_103878007.1 (DdbA), WP_008767132.1 (DdbB), WP_008762511.1 (DdbC), WP_048698952.1 (DdbD), WP_146072611.1 (DdbE), and WP_103878006.1 (DdbF). Putative signal peptides in protein amino acid sequences were identified using SignalP 5.0 (37). Lipoprotein export signals were identified based on presence of a previously described C(x/xx)(D/E)(D/E) amino acid sequence (with x signifying “any amino acid” and the first cystine corresponding to the predicted lipidation site) (21–23).

To investigate the evolutionary distribution and diversity of *ddbABCDEF*-like loci, we conducted comparative genomic analyses across gut Bacteroidales using publicly available genomes from NCBI RefSeq. Distinct subtypes of *ddbABCDEF*-like loci were identified in *B. thetaiotaomicron* and *P. vulgatus*, designated as Bt1 (*B. thetaiotaomicron* DFI.1.47, NZ_JAJCPU010000061:25232..34119) and Bt2 (*B. thetaiotaomicron* MSK.14.51, NZ_JAHOLZ010000003:complement(83009..91766)), as well as Pv1 (*P. vulgatus* MSK.23.57, NZ_JAHOBM010000001:10853..115503) and Pv2 (*P. vulgatus* MSK.9.20, NZ_JAHPXN010000009:50891..56657). Additionally, we identified a homologous locus in *Riemerella anatipestifer* ATCC 11845 (NC_017045:1014432..1020133).

To determine the taxonomic distribution of *ddb* subtypes, protein sequences of DdbC homologs from *B. thetaiotaomicron* (Bt1: WP_008762511.1, Bt2: WP_055220680.1) and *P. vulgatus* (Pv1: WP_005843284.1, Pv2: WP_117829583.1) were used as queries (**Table S3**) for BLASTp searches against a database comprising the proteomes of 9,910 Bacteroidota genomes with unambiguous genus and species designations downloaded from NCBI (**Table S4**). Blastp hits with an e-value ≤ 1e-15, with >50% sequence identity and >75% sequence coverage to the query sequences were considered significant.

## ACKNOWLEDGEMENTS

Research reported in this publication was supported by funding from the National Institutes of Health (NIGMS R35GM146969), the Searle Scholars Program, and the Duchossois Family Institute (to S.H.L). We thank Dr. Eric Pamer and the Duchossois Family Institute Microbiome Metagenomics and Host-Microbe Metabolomics Facilities for experimental support.

---

**Figure S1.**
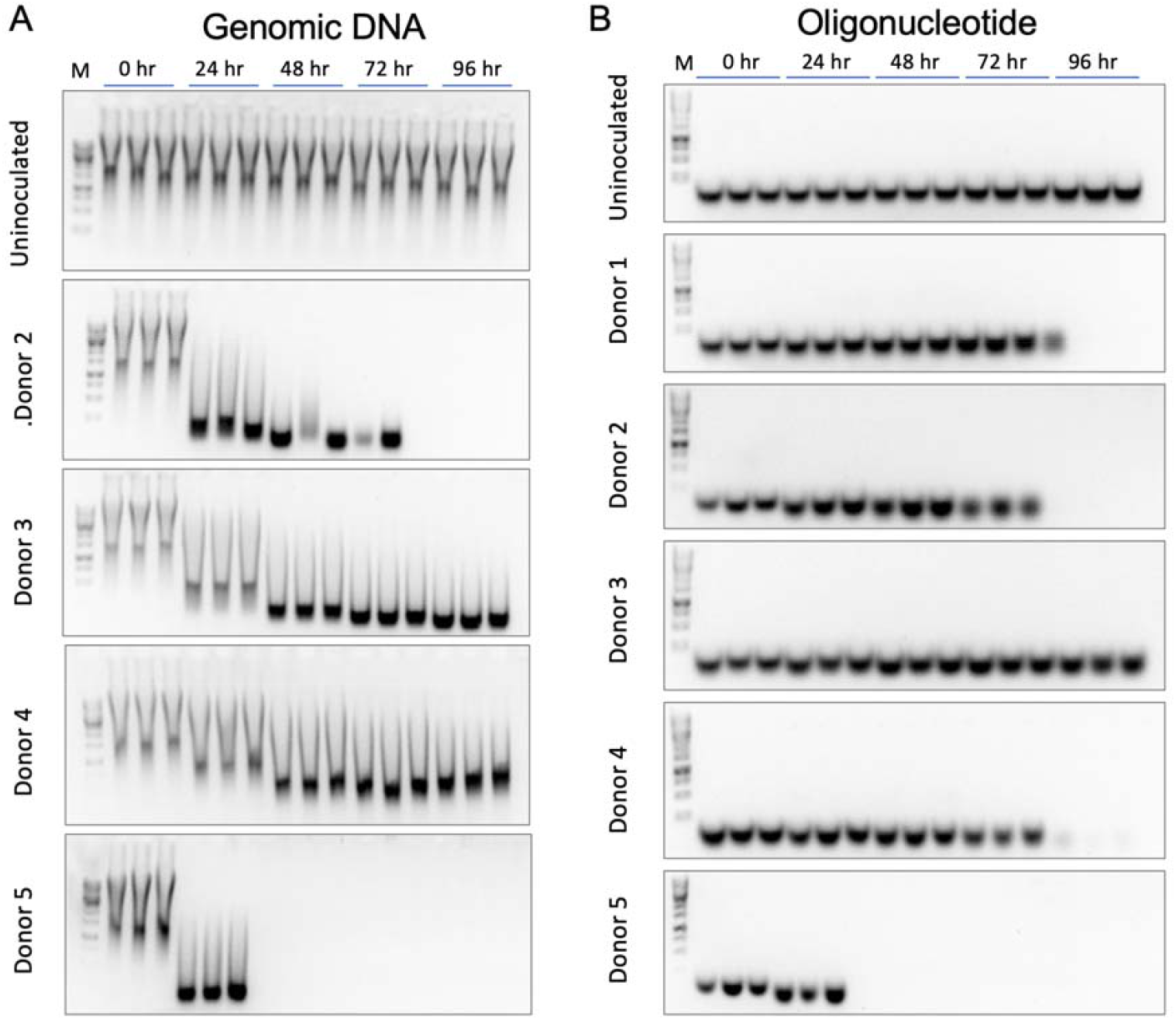
DNA-depleting activity of human fecal microbial communities. Agarose gels showing progressive degradation of (A) herring sperm DNA and (B) Tracking of arbitrary sequence 60-base pair double-stranded oligonucleotides incubated with fecal slurries from distinct donors.

**Figure S2.**
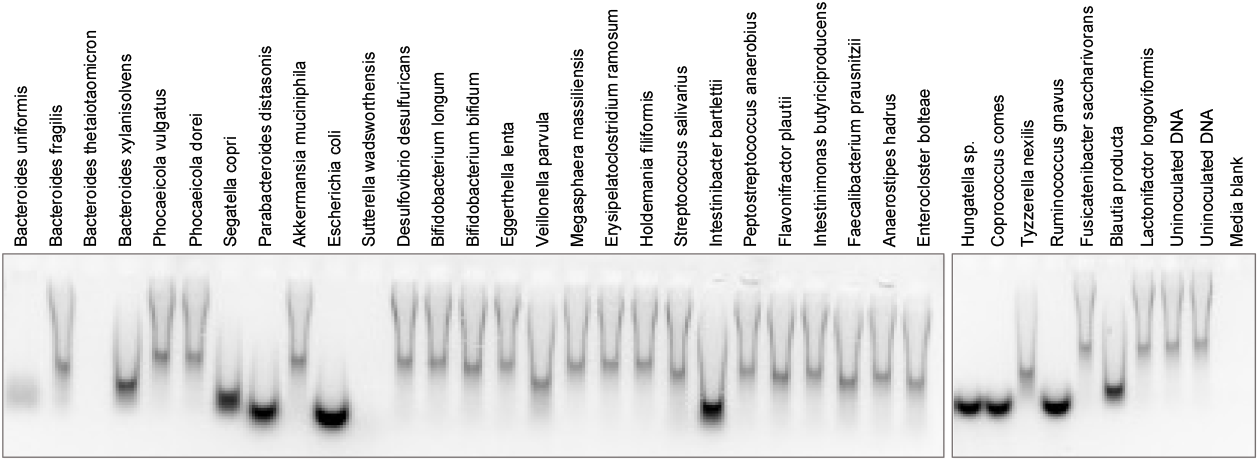
DNA Depletion activity of Individual strains. Agarose gel images showing DNA degradation by human gut bacterial isolates.

**Figure S3.**
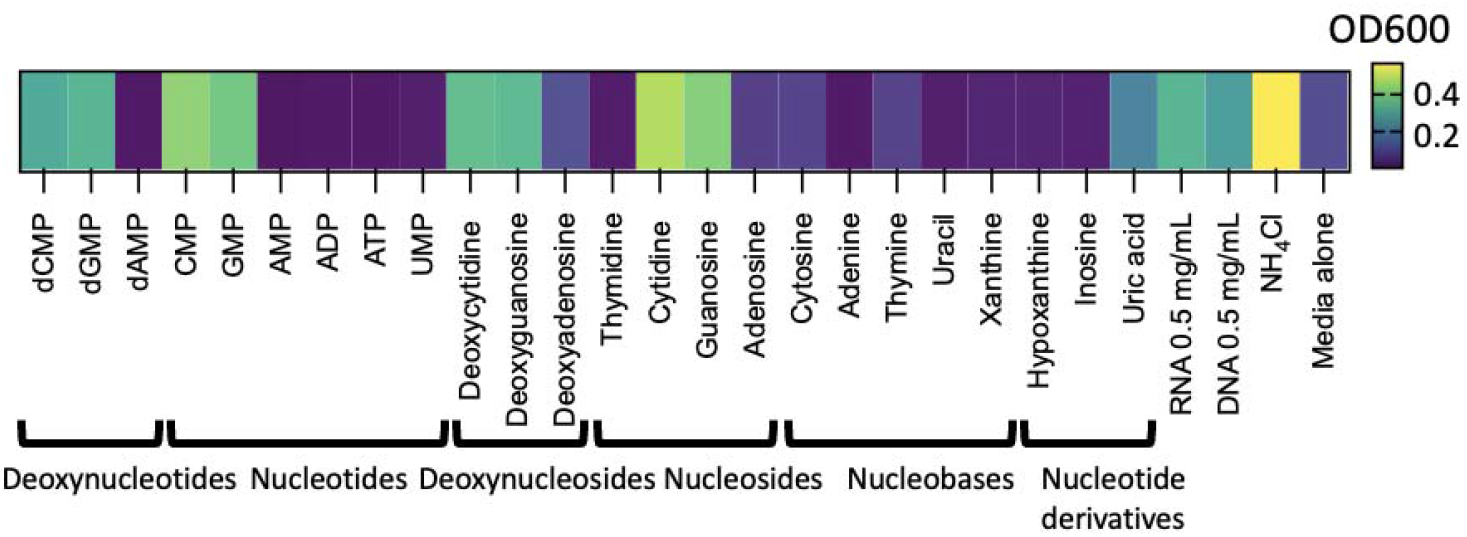
Maximal growth of *B. thetaiotaomicron* DFI.1.47 in nitrogen-free minimal media supplemented with different potential nitrogen sources. Growth in chemically defined “nitrogen-free” medium supplemented with indicated compounds. Three biological replicates were tested for each condition, with values representing the average of these replicates.

**Figure S4.**
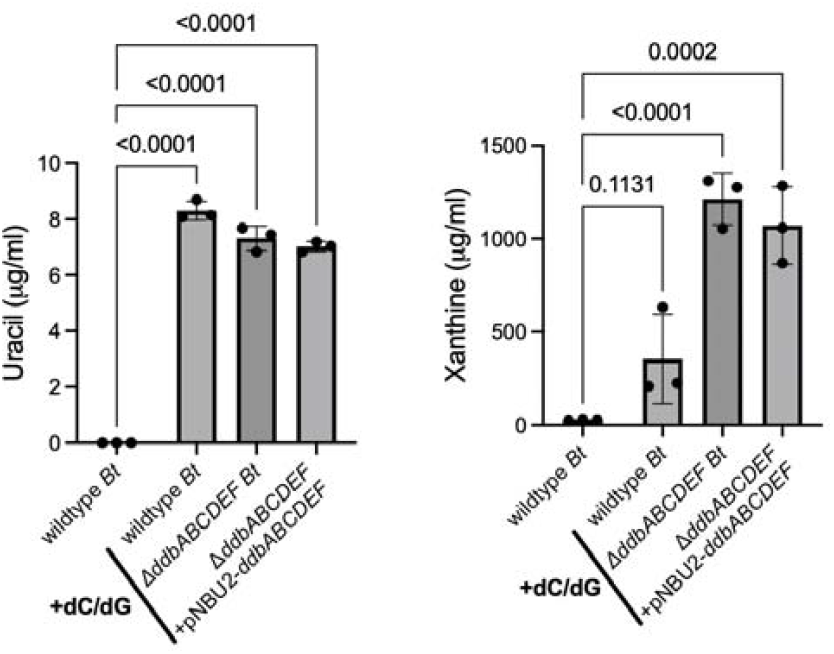
*ddbABCDEF* is dispensable for nucleobase production from nucleosides. . Quantification of uracil and xanthine levels in culture supernatants of *B. thetaiotaomicron* grown with or without exogenous deoxycytidine and deoxyguanosine.

## Notes

### Competing Interest Statement

The authors have declared no competing interest.

